# Sphingosine Kinase 2 associates with the nsP3 of chikungunya virus and is required for replication

**DOI:** 10.1101/2020.09.10.291682

**Authors:** Opeoluwa O. Oyewole, St Patrick Reid

## Abstract

Sphingosine kinase 2 (SK2) is a lipid kinase that catalyzes the production of sphingosine-1-phosphate (S1P) from sphingosine. Previously, we have shown that SK2 is recruited to the viral replication complex (VRC) early during chikungunya virus (CHIKV) infection. In the present study, we demonstrate that SK2 is required for viral replication and protein production. Treatment with a SK2 inhibitor significantly impaired the function of a CHIKV replicon. Similarly, compound treatment or genetic targeting resulted in impaired viral protein production. Mechanistically, we demonstrate that CHIKV nsP3 binds to SK2. Association of nsP3 with SK2 was mediated, in part, through the FGDF motifs within the hypervariable domain (HVD) of nsP3. In a competition assay, SK2 competed with G3BP for binding to nsP3. Collectively, these results extend our previous findings and identify SK2 as a CHIKV host factor recruited by nsP3.

## 1. Introduction

Chikungunya virus (CHIKV) is a re-emerging mosquito-borne enveloped alphavirus in the *Togaviridae* family. The virus is the etiologic agent of Chikungunya fever (CHIKF), a disease characterized by high fever and debilitating polyarthralgia and polyarthritis that can typically last for 1-4 weeks (Simon, Javelle et al. 2011). In some cases, individuals develop chronic arthritis that lasts for months to years following infection (Kennedy, Fleming et al. 1980, Pialoux, Gauzere et al. 2007, Borgherini, Poubeau et al. 2008, Schilte, Staikowsky et al. 2013). Although CHIKV infection rarely results in mortality, the chronic stage of the disease can lead to a high economic burden during outbreaks (Cardona-Ospina, Villamil-Gomez et al. 2015). Currently, there are no approved vaccines and therapeutics available to prevent or treat CHIKV infection.

CHIKV has a positive-strand RNA genome that encodes four nonstructural proteins (nsP1-nsP4) and five structural proteins (capsid, E3, E2, 6K, and E1) from two open reading frames. Upon entry, the newly synthesized nsPs drive replication and transcription of the viral RNA. Of the nsPs, nsP1, nsP2, and nsP4 all possess well-characterized enzymatic activities that are essential for replication (Kaariainen and Ahola 2002). The precise function of nsP3, however, has remained elusive, although studies evaluating the function of nsP3 mutations indicate a role in viral RNA synthesis (LaStarza, Lemm et al. 1994, De, Fata-Hartley et al. 2003). A recent study suggested nsP3 possesses hydrolase activity, which is required for optimal viral replication (Abraham, Hauer et al. 2018).

CHIKV nsP3, similar to other alphavirus nsP3s, is a modular protein with three distinct domains. The N-terminus of nsP3 contains a highly conserved macrodomain (Malet, Coutard et al. 2009, Neuvonen and Ahola 2009). Recent studies have shown that the macrodomain has mono-ADP-ribosylhydrolase activity that is critical for replication and virulence (Eckei, Krieg et al. 2017, McPherson, Abraham et al. 2017, Abraham, Hauer et al. 2018). The central region of nsP3 binds Zn^2+^, and, although its function in viral replication is unclear, Zn^2+^ binding mutants failed to express the polyprotein and replicate virus, indicating a critical role during infection (Shin, Yost et al. 2012). The C-terminal domain of nsP3 is highly phosphorylated and disordered. Interestingly, this domain shows very little sequence conservation between alphaviruses and as result has been termed the hypervariable domain (HVD). This domain has been demonstrated to act as an assembly hub for recruiting host factors to the viral replication complex (VRC) (Foy, Akhrymuk et al. 2013, Foy, Akhrymuk et al. 2013, Kim, Reynaud et al. 2016, Frolov, Kim et al. 2017). Importantly, the HVD of different alphaviruses appear to recruit distinct sets of host proteins to the VRC during infection (Foy, Akhrymuk et al. 2013, Frolov, Kim et al. 2017). Host proteins including, G3BP, FXR, Amphiphysins, CD2AP, NAP proteins (NAP1L1 & NAP1L4), and SH3KBP1 have all been shown to be recruited by nsP3 via the HVD (Neuvonen, Kazlauskas et al. 2011, Fros, Domeradzka et al. 2012, Foy, Akhrymuk et al. 2013, Frolov, Kim et al. 2017, Meshram, Agback et al. 2018, Mutso, Morro et al. 2018).

We have recently shown that the lipid kinase, sphingosine kinase 2 (SK2), is an essential host factor during CHIKV infection. Knockdown of SK2 or pre-treatment with small molecule inhibitors significantly inhibited virus infection. SK2 was demonstrated to co-localize with the viral RNA and viral proteins associated with the VRC early during infection, suggesting it plays a role in viral replication. Additionally, a GFP-tagged nsP3 co-localized with SK2 (Reid, Tritsch et al. 2015). In the current study, we extend these findings and demonstrate that SK2 is required for efficient replication and production of viral proteins. Further, CHIKV nsP3 interacted with SK2 during infection. Domain mapping studies revealed that SK2 associates with the HVD of nsP3 and can compete with G3BP for nsP3 binding. Taken together, we have identified the viral factor, nsP3, necessary for the recruitment of the novel CHIKV host factor SK2 to the VRC.

## 2. Results

### 2.1. SK2 is required for optimal replication and viral protein production

Previously, we identified SK2 as an essential CHIKV host factor (Reid, Tritsch et al. 2015). Co-localization and proteomic experiments suggested a role for the kinase in the VRC. To explore a role for the kinase in replication, we used the BHK-CHIKV-NCT replicon cell line. The cell line stably expresses the CHIKV nsPs which drive the expression of EGFP and Renilla luciferase markers (Pohjala, Utt et al. 2011). Treatment of the cells with increasing doses of the well-characterized SK2-specific inhibitor ABC294640 (ABC) resulted in significant reduction of GFP expression (Fig. 1A). In our previous study, 30 μM of ABC severely impaired viral infection; similarly, here 30 μM inhibited replicon activity by ∼80%. Ribavirin, previously shown to inhibit replicon activity was used as a positive control and displayed greater than 80% inhibitory activity in these experiments (Fig. 1A). Representative images collected on a high content imager are shown in Figure 1B. We next explored early protein production during infection. Cells were pretreated with either 3 or 30 μM of ABC or vehicle, and then infected. Viral protein production was monitored 2, 4, and 6 h post infection. Pretreatment with 30 μM of ABC significantly impaired early viral protein production at 4 h and 6 h compared to untreated and 3 μM pretreated samples (Fig. 1C). Next, we evaluated early viral protein production after CRISPR-mediated knockout of SK2 expression. Similar to ABC treatment, knockout of SK2 resulted in a clear impairment of early viral protein production (Fig. 1D). Taken together, these results demonstrate that targeting SK2 impairs CHIKV replication and early viral protein production, suggesting a role for the kinase in viral replication.

**Figure 1.**
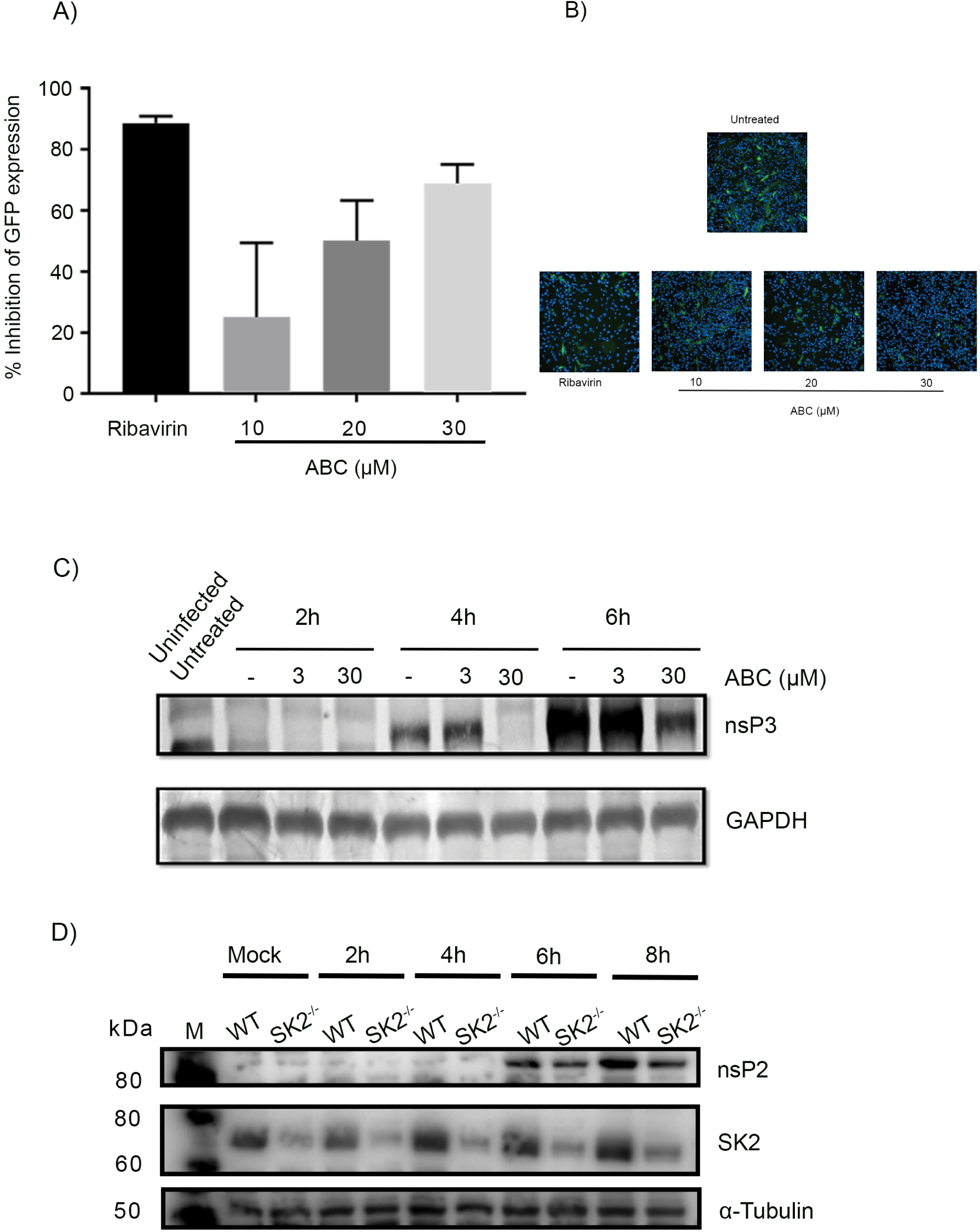
SK2 is required for CHIKV replication and viral protein production. (A) BHK-CHIKV-NCT cells were treated with 30 μM ribavirin or with 10, 20, or 30 μM ABC294640 (ABC). Twenty-four hours post-treatment, the cells were fixed and analyzed for EGFP expression by high content confocal microscopy. Values represent mean + SEM (n=6). (B) Representative images showing GFP expression of treated cells in (A) (Scale bar = 100 µm. (C) BHK21 cells were pretreated with 3 or 30 μM ABC then infected with CHIKV (MOI = 5) for the indicated time points. Cell lysates were harvested and subjected to western blot analysis to detect nsP3 and GAPDH. (D) SK2^-/-^ HeLa cells were mock-infected or infected with CHIKV (MOI = 5) for the indicated time points. Cell lysates were harvested and subjected to western blot analysis of SK2, nsP2, and GAPDH expression.

### 2.2. SK2 interacts with the nsP3

Expression of an EGFP-tagged nsP3 alone resulted in recruitment of SK2 from the nucleus to distinct nsP3-positive puncta, suggesting an association of nsP3 with SK2 (Reid, Tritsch et al. 2015). To test if nsP3 specifically interacts with SK2 during infection, we performed co-immunoprecipitation (co-IP) experiments. Cells were infected with CHIKV for 24 h, and the lysates subjected to IP using an anti-SK2 antibody or Ig control. Here, nsP3 expressed during infection was found to interact with SK2 and not with the control (Fig 2A), thus demonstrating a specific interaction of nsP3 with SK2 during infection.

**Figure 2.**
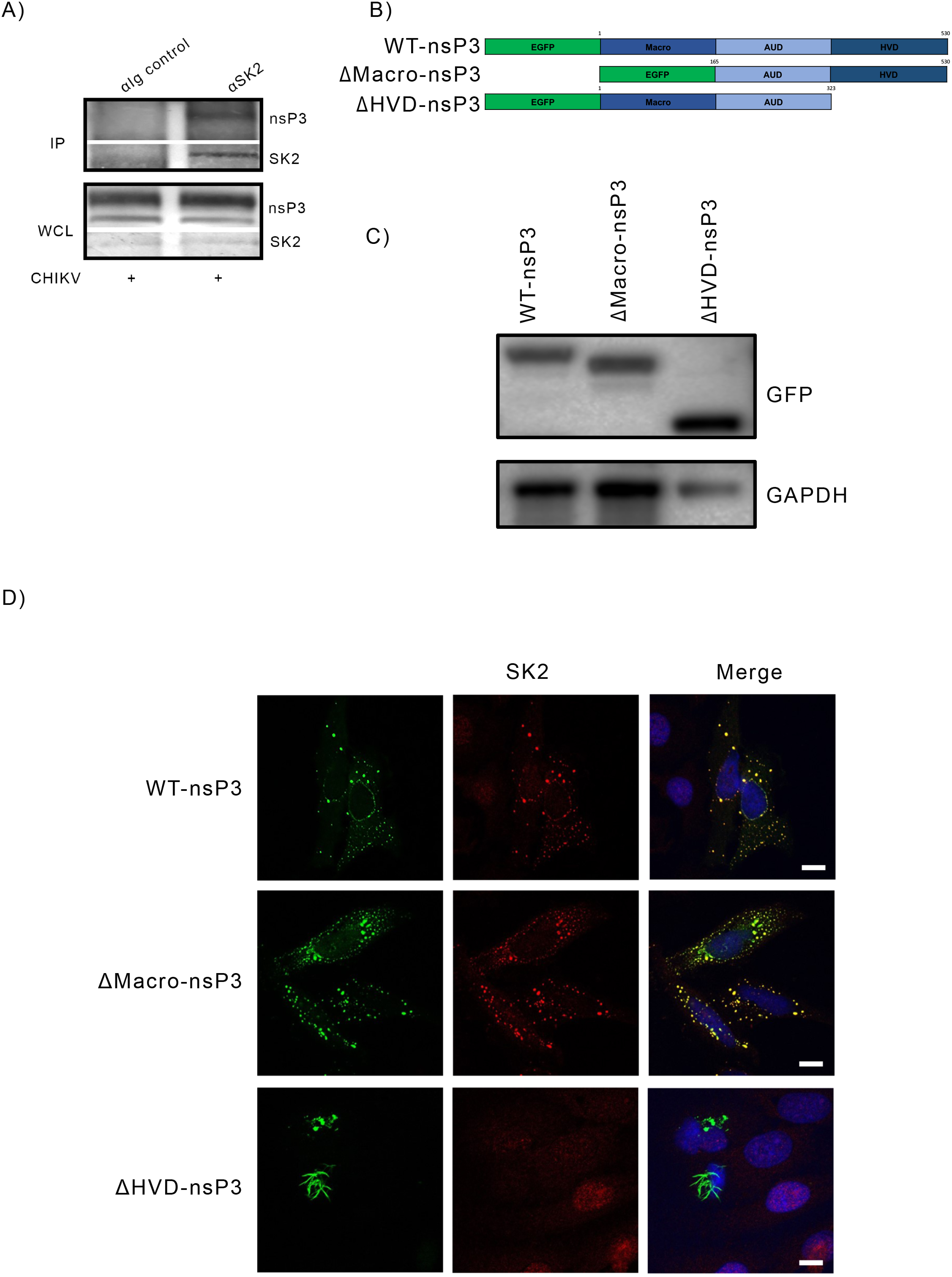
The Hypervariable domain (HVD) of CHIKV is essential for co-localization with SK2. (A) BHK21 cells were infected with CHIKV at a MOI of 5 for 24 h. The cell lysates were then subjected to immunoprecipitation (IP) with either SK2 polyclonal antibody or Ig control antibody followed by western blot analysis to detect nsP3 and SK2. Whole cell lysate (WCL) was also analyzed to confirm expression of nsP3 and SK2. (B) Schematic representation of the WT-nsP3 and the deletion mutants. (C and D) HeLa cells were transfected with WT-nsP3, nsP3-ΔMacro, and nsP3-ΔHVD and (C) protein expression was confirmed by western blot using anti-GFP and anti-GAPDH antibody. (D) EGFP-fused nsP3 constructs (green), SK2 (red), and Hoechst nuclear stain (blue) were visualized by confocal microscopy. Scale bar = 5 μm.

### 2.3. The Hypervariable domain of nsP3 is required for SK2 recruitment

The HVD of nsP3 has been shown to interact with host cell factors during infection. We constructed a series of EGFP-tagged mutants in which the macro domain or the HVD was deleted and expression confirmed by western blot (Fig 2B & 2C). Next, we performed immunofluorescence (IF) experiments to determine SK2 localization in the presence of either WT nsP3 or the nsP3 truncations. Twenty-four hours post transfection, the cells were analyzed for endogenous SK2 and EGFP localization. WT nsP3 induced a dramatic re-localization, similar to previous studies (Reid, Tritsch et al. 2015). Similarly, deletion of the macrodomain induced a re-localization of SK2. Deletion of the HVD failed to re-localize SK2 (Fig 2D). Interestingly, the HVD deletion mutant localization was dramatically altered, which could explain the lack of interaction. It should be noted that the localization seen here is similar to prior studies in which HVD deletion dramatically affected nsP3 localization (Fros, Domeradzka et al. 2012). These data suggest the HVD is required for SK2-nsP3 co-localization.

### 2.4. The nsP3 FGDF motif is required for SK2 interaction

The HVD acts as a hub for cellular protein recruitment. Within the HVD domain, specific motifs have been identified that are required for specific host-protein interactions. For example, the FGDF repeat motifs interact with G3BP proteins and the proline-rich region interacts with Amphiphysins (Neuvonen, Kazlauskas et al. 2011, Kim, Reynaud et al. 2016). To identify the region of the HVD required for SK2 interaction we constructed deletion mutants of known nsP3 regions. Truncation mutations in which the FGDF motifs were removed lost the ability to co-localize with SK2 (data not shown). To investigate the involvement of the FGDF motifs, we generated, and confirmed expression of, single and double alanine substitution mutants in which either the N-terminal FGDF motif (FGDF_N_Ala), the C-terminal FGDF motif (FGDFcAla), or both motifs (FGDF_N+C_Ala) were mutated (Fig 3A & 3B). Each mutant was tested for co-localization with SK2 via IF. The FGDFcAla single substitution mutant retained the ability to co-localize with SK2 similar to WT. However, co-localization with the kinase was lost in the double substitution mutant (Fig 3C). We confirmed this interaction via co-Ip and observed that interaction was lost in the FGDF_N_Ala mutant and the double mutant (FGDF_N+C_Ala) (Fig 3D). G3BP has previously been reported to be associated with nsP3 FGDF motifs (Kim, Reynaud et al. 2016). Therefore, to further test association of SK2 with the FGDF motif we performed a competition assay in which an HA-tagged nsP3 (nsP3-HA) was used to immunoprecipitate G3BP1 in the presence and absence of a Flag-tagged SK2 (Flag-SK2). In the absence of Flag-SK2, we observed G3BP association with nsP3-HA. However, when increased amounts of Flag-SK2 were introduced, nsP3-G3BP1 was lost and a concomitant increase in nsP3-SK2 association was observed (Fig. 3E). Our results demonstrate that the interaction between nsP3 and SK2 is mediated in part by the FGDF motifs and that SK2 can compete with G3BP1 for nsP3 interaction.

**Figure 3.**
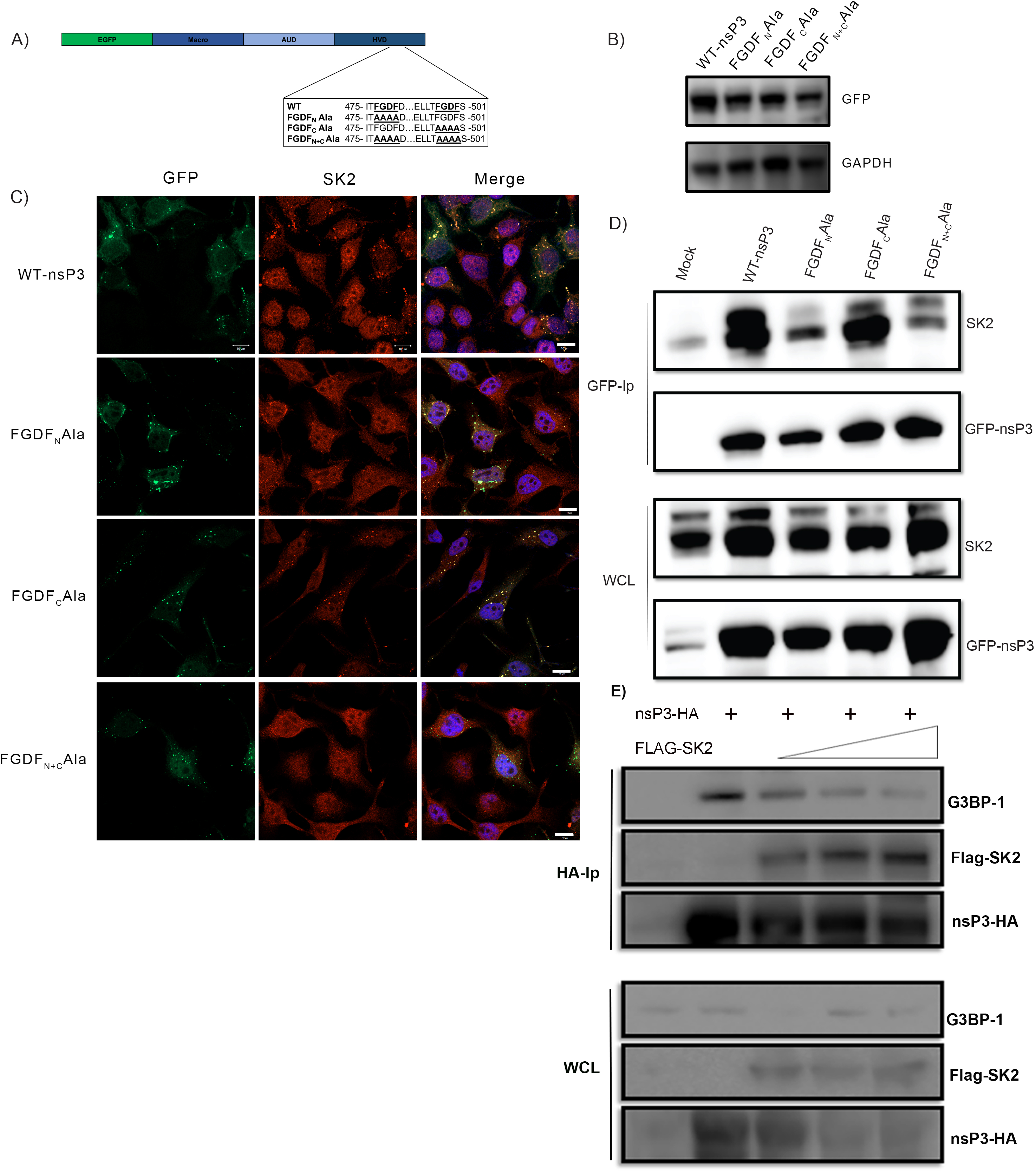
FGDF motifs in CHIKV nsP3-HVD are crucial in recruitment of SK2. (A) Schematic showing alanine substitutions of the nsP3 HVD. (B and C) HeLa cells were transfected with EGFP-nsP3 constructs containing WT or alanine substitution FGDF mutations - FGDF_N_Ala-nsP3, FGDF_C_Ala-nsP3, FGDF_N+C_Ala-nsP3 and 24 h post transfection, (B) cells were lysed and protein expression was confirmed by western blot and (C) cells were fixed and stained for EGFP-fused nsP3 (green) SK2 (red), and Hoechst nuclear stain (blue) expression and visualized by confocal microscopy. Scale bar = 10 μm. (D) HeLa cells were transfected with EGFP-nsP3 constructs containing alanine substitution mutations - FGDF_N_Ala-nsP3, FGDF_C_Ala-nsP3, FGDF_N+C_Ala-nsP3. 24 h post transfection, cells lysates were harvested and subjected to immunoprecipitation using GFP-conjugated beads followed by western blot to detect endogenous SK2 and GFP. Whole cell lysate was also analyzed to confirm expression of SK2 and GFP-nsP3 (E) HeLa cells were transfected with nsP3-HA and increasing concentration of Flag-SK2 (0.5, 1, and 2 μg) where indicated. Twenty-four hours post-transfection, cell lysates were harvested and subjected to immunoprecipitation using HA-beads followed by western blot analysis to detect endogenous G3BP1, Flag-SK2, and nsP3-HA. Whole cell lysate (WCL) was also analyzed to confirm expression of G3BP1, nsP3-HA and Flag-SK2.

### 2.5. Concluding remarks

While the precise function(s) of the various proteins recruited by the HVD of nsP3 have yet to be fully elucidated, it is increasingly evident that they are critical for viral replication (Lark, Keck et al. 2017, Gotte, Liu et al. 2018). Similarly, we postulate that the lipid kinase SK2 is recruited to the VRC for a role in viral replication. Several lines of evidence suggest SK2 indeed plays a novel role in viral replication. First, in our previous study, SK2 associated with viral dsRNA in the VRC, and proteomic analysis revealed an association of SK2 with the viral polyprotein and host proteins associated with mRNA processing and transcription during infection (Reid, Tritsch et al. 2015). Here, we report that impairment of SK2 function significantly inhibited CHIKV replication in a replicon-based system (Fig 1A). Additionally, impairment of SK2 function or reduced expression of the kinase affected viral protein production, indicating the kinase is required for optimal viral replication.

SK2 along with SK1 phosphorylates sphingosine to the bioactive lipid, S1P (Adams, Pyne et al. 2016). As key regulators of sphingolipid metabolism, these kinases play a pivotal role in a number of cellular and physiological processes. SK1, the more extensively studied isozyme, has been demonstrated to be a key modulator during infection of numerous viruses (Carr, Mahalingam et al. 2013). Recent studies have shown that, similar to SK1, SK2 also plays a role during viral infection. The first such study demonstrated that SK2 enhances KSHV latency, while a second study showed that lipid peroxidation, regulated in part by SK2, could restrict HCV replication (Dai, Plaisance-Bonstaff et al. 2014, Yamane, McGivern et al. 2014). More recently, using an apoptosis-related siRNA library screen, SK2 was demonstrated to play a pro-apoptotic role in DENV-infected hepatic cells (Morchang, Lee et al. 2017). In a similar manner, we identified a role for SK2 in CHIKV infection from a human kinome siRNA library screen. Although we demonstrated that SK2 was critical for CHIKV infection, the mechanism of SK2 recruitment was unclear (Reid, Tritsch et al. 2015).

In the present study, we demonstrate that SK2 plays a role in viral replication at least in part through an association with the FGDF motifs in the HVD of nsP3. Similar to SK2, G3BP is recruited to the VRC through an association with the FGDF motifs in the HVD of nsP3. Early G3BP binding with the CHIKV VRC is a prerequisite for RNA replication (Kim, Reynaud et al. 2016). This function requires domains of G3BPs involved in protein-protein interaction and RNA-binding (Kim, Reynaud et al. 2016). While it is currently unclear whether SK2 possesses similar functional domains, SK2 has been shown to modulate DNA synthesis (Igarashi, Okada et al. 2003, Kihara, Anada et al. 2006). Further investigation into these potential functional properties of SK2 will be required to delineate if SK2 associates with the VRC in a similar manner to G3BP.

A competition assay revealed that SK2 could compete with G3BP1 for binding to nsP3. This observation confirms the requirement of the FGDF motif for SK2 binding. However, it is currently unclear how this observation affects nsP3 VRC protein-protein interaction dynamics. For example, are SK2-nsP3 and G3BPs-nsP3 in distinct complexes within the VRC? Further studies will be required to delineate the dynamics of these interactions. Moreover, does G3BP associate with SK2? In the absence of CHIKV infection, SK2 is primarily localized in the nucleus whereas G3BPs are typically found diffuse in the cytoplasm. IF analysis of both proteins in the absence of nsP3 or viral infection did not show co-localization (unpublished observations).

SK2 along with SK1 are targets for anti-cancer treatments (Shida, Takabe et al. 2008, Hasanifard, Sheervalilou et al. 2019). As a result, a number of potent inhibitors have been developed against these kinases. Notably, the compound ABC used in the present study, displays potent anti-tumor activities against a variety of cancers (French, Upson et al. 2006, Beljanski, Knaak et al. 2010). The compound is currently under evaluation in a phase I clinical trial for patients with solid tumors (Clinical trials.gov identifier: NCT01488513) and in a phase I/II clinical trial for HIV+ patients with diffuse large B-cell lymphoma (Clinicaltrials.gov identifier: NCT02229981). Our current study, along with the prior study, suggests targeting SK2 can serve as a viable host-based targeting approach to inhibiting viral replication. Future studies examining SK2 inhibitors will provide greater insight into the feasibility of this approach. Nevertheless, the requirement of SK2 for CHIKV infection and the availability of potent SK2 inhibitors provides a clear rationale moving forward.

## 3. Materials and Methods

### 3.1. Cell Culture, Virus and Reagents

Baby Hamster Kidney (BHK)-21 and HeLa cells were purchased from American Type Culture Collection (ATCC; Manassas, VA, USA). The cells were cultured in Dulbecco’s modified Eagle’s medium (DMEM) (Life Technologies, Carlsbad, CA, USA) supplemented with 10% inactivated fetal bovine serum (FBS) (GE Healthcare, Piscataway, NJ, USA). ABC294640 was purchased from Cayman Chemical (Ann Arbor, MI, USA) and Ribavirin was purchased from Sigma (St. Louis, MO, USA).

CHIKV (CHIKV-181/25), was propagated from the USAMRIID collection of viruses. The CHIKV EGFP-nsP3 construct was used as previously reported (Reid, Tritsch et al. 2015). The EGFP-nsP3 truncations and alanine substitution mutations were constructed by subcloning the corresponding nsP3-encoding PCR products into the vector pEGFP-C1 (Clontech) using XhoI-KpnI restriction sites.

### 3.2. Western Blot Experiments

For the viral co-immunoprecipitation experiments, BHK21 cells were infected with CHIKV at a MOI of 5 for 1 h in serum-free DMEM. After adsorption, the cells were washed once, and complete medium was added. Twenty-four hours post infection, the cells were lysed in lysis buffer (50mM Tris (pH 7.4), 150 mM NaCl, 1% NP-40, 0.25% Na-deoxycholate, 1 mM ethylenediaminetetraacetic acid, protease inhibitor (Complete: Roche, Nutley, NJ, USA), and PhosSTOP phosphatase inhibitor (Roche, Nutley, NJ, USA)) and immunoprecipitated using anti-SK2 antibody or an Ig control antibody (ECM Biosciences, Versailles, KY, USA). After a 1 h antibody incubation, protein complexes were precipitated using Protein A agarose beads (Sigma, St. Louis, MO, USA). The precipitated material was analyzed by western blot, using the anti-nsP3 antibody (kindly provided by Dr. Andres Merits) and the aforementioned anti-SK2 antibody.

For detection of truncation and substitution mutants, 2 µg of each plasmid was transfected into HeLa cells in a 6-well plate. After 24 hours, the cells were lysed in lysis buffer and protein expression was analyzed by Western blot, using rabbit anti-GFP (Sigma, St. Louis, MO, USA) and rabbit anti-GAPDH (Sigma, St Louis, MO, USA) antibodies.

For the competition co-immunoprecipitation experiments, HeLa cells were transfected with nsP3-HA in the absence or presence of increasing concentration of Flag-SK2 (0.5, 1 and 2 μg) using the JetPRIME Polyplus transfection kit (Polyplus Transfection, New York, NY, USA). Twenty-four hours post-transfection, cells were lysed in lysis buffer and precipitated using a monoclonal HA antibody cross-linked to agarose beads (Sigma, St Louis, MO, USA). The precipitated material was analyzed by western blot. Endogenous G3BP was detected with a mouse monoclonal antibody (BD Biosciences, San Jose, CA, USA). Flag and HA were detected with mouse monoclonal antibodies (Invitrogen, Carlsbad, CA, USA).

### 3.3 Detection of CHIKV replication via EGFP expression

Confluent BHK-CHIKV-NCT cells grown in 96-well plates were treated with Ribavirin (30 μM) or various concentrations of ABC (10, 20 and 30 μM). Twenty-four hours post treatment, the cells were fixed in 4% PFA, and subsequently, cell nuclei and cytoplasm labeled with Hoechst 33342 (Life Technologies, Carlsbad, CA, USA) and HCS CellMask Red (Life Technologies, Carlsbad, CA, USA), respectively. High content quantitative imaging data were acquired and analyzed on an Operetta CLS high content system (Perkin Elmer, Boston, MA, USA). Image analysis was accomplished using standard Acapella scripts. Percentage inhibition of EGFP expression was calculated using the mean percentage of EGFP expression from untreated cells.

### 3.4 CRISPR-mediated SK2 knockout

HeLa cells were transfected with a GFP-expressing plasmid containing guideRNAs and Cas9 targeting SK2 (1 µg) and an RFP-expressing plasmid containing the HDR and puromycin selection marker purchased from SantaCruz Biotechnology (Dallas, TX USA), using JetPRIME Polyplus transfection kit. Seventy-two hours post-transfection, transfected cells were selected using Puromycin (2 µg/mL) for 2 weeks. The cells were subsequently sorted via fluorescence-activated cell sorting (FACS) for cells expressing both GFP and RFP and analyzed for protein expression by western blot using the aforementioned anti-SK2 antibody.

For the time-course study, WT or SK2 knockout cells were infected with CHIKV (MOI = 5) for 2, 4, 6, or 8 hours. The cells were subsequently lysed and analyzed for protein expression by western blot using the anti-nsP2 antibody (kindly provided by Dr. Andres Merits) and the aforementioned anti-SK2 antibody.

### 3.5. Immunofluorescence Experiments

Transfected HeLa cells were fixed in 4% paraformaldehyde, blocked in PBS-3%BSA for 1 h and then stained with the indicated antibodies. SK2 was detected with rabbit anti-SK2. Secondary Alexa 568-conjugated goat anti-rabbit antibody (Life Technologies, Carlsbad, CA, USA) was used to visualize SK2 primary antibody. Expression of nsP3 plasmids was observed as GFP expression by confocal microscopy. Cell nuclei were labeled with Hoechst 33342 (Life Technologies, Carlsbad, CA USA). Images were collected using the Zeiss LSM800 confocal microscope.

## Acknowledgements

We thank Janice A. Taylor and James R. Talaska of the Advanced Microscopy Core Facility at the University of Nebraska Medical Center for providing assistance with confocal microscopy. We also thank Dr. Scott Ouellette for critical review of the manuscript. This work was supported by startup funds for SPR.

## Notes

### Competing Interest Statement

The authors have declared no competing interest.

## References

Abraham, R., D. Hauer, R. L. McPherson, A. Utt, I. T. Kirby, M. S. Cohen, A. Merits, A. K. L. Leung and D. E. Griffin (2018). “ADP-ribosyl-binding and hydrolase activities of the alphavirus nsP3 macrodomain are critical for initiation of virus replication.” Proc Natl Acad Sci U S A 115(44): E10457–E10466.

Adams, D. R., S. Pyne and N. J. Pyne (2016). “Sphingosine Kinases: Emerging Structure-Function Insights.” Trends Biochem Sci 41(5): 395–409.

Beljanski, V., C. Knaak and C. D. Smith (2010). “A novel sphingosine kinase inhibitor induces autophagy in tumor cells.” J Pharmacol Exp Ther 333(2): 454–464.

Borgherini, G., P. Poubeau, A. Jossaume, A. Gouix, L. Cotte, A. Michault, C. Arvin-Berod and F. Paganin (2008). “Persistent arthralgia associated with chikungunya virus: a study of 88 adult patients on reunion island.” Clin Infect Dis 47(4): 469–475.

Cardona-Ospina, J. A., W. E. Villamil-Gomez, C. E. Jimenez-Canizales, D. M. Castaneda-Hernandez and A. J. Rodriguez-Morales (2015). “Estimating the burden of disease and the economic cost attributable to chikungunya, Colombia, 2014.” Trans R Soc Trop Med Hyg 109(12): 793–802.

Carr, J. M., S. Mahalingam, C. S. Bonder and S. M. Pitson (2013). “Sphingosine kinase 1 in viral infections.” Rev Med Virol 23(2): 73–84.

Dai, L., K. Plaisance-Bonstaff, C. Voelkel-Johnson, C. D. Smith, B. Ogretmen, Z. Qin and C. Parsons (2014). “Sphingosine kinase-2 maintains viral latency and survival for KSHV-infected endothelial cells.” PLoS One 9(7): e102314.

De, I., C. Fata-Hartley, S. G. Sawicki and D. L. Sawicki (2003). “Functional analysis of nsP3 phosphoprotein mutants of Sindbis virus.” J Virol 77(24): 13106–13116.

Eckei, L., S. Krieg, M. Butepage, A. Lehmann, A. Gross, B. Lippok, A. R. Grimm, B. M. Kummerer, G. Rossetti, B. Luscher and P. Verheugd (2017). “The conserved macrodomains of the non-structural proteins of Chikungunya virus and other pathogenic positive strand RNA viruses function as mono-ADP-ribosylhydrolases.” Sci Rep 7: 41746.

Foy, N. J., M. Akhrymuk, I. Akhrymuk, S. Atasheva, A. Bopda-Waffo, I. Frolov and E. I. Frolova (2013). “Hypervariable domains of nsP3 proteins of New World and Old World alphaviruses mediate formation of distinct, virus-specific protein complexes.” J Virol 87(4): 1997–2010.

Foy, N. J., M. Akhrymuk, A. V. Shustov, E. I. Frolova and I. Frolov (2013). “Hypervariable domain of nonstructural protein nsP3 of Venezuelan equine encephalitis virus determines cell-specific mode of virus replication.” J Virol 87(13): 7569–7584.

French, K. J., J. J. Upson, S. N. Keller, Y. Zhuang, J. K. Yun and C. D. Smith (2006). “Antitumor activity of sphingosine kinase inhibitors.” J Pharmacol Exp Ther 318(2): 596–603.

Frolov, I., D. Y. Kim, M. Akhrymuk, J. A. Mobley and E. I. Frolova (2017). “Hypervariable Domain of Eastern Equine Encephalitis Virus nsP3 Redundantly Utilizes Multiple Cellular Proteins for Replication Complex Assembly.” J Virol 91(14).

Fros, J. J., N. E. Domeradzka, J. Baggen, C. Geertsema, J. Flipse, J. M. Vlak and G. P. Pijlman (2012). “Chikungunya virus nsP3 blocks stress granule assembly by recruitment of G3BP into cytoplasmic foci.” J Virol 86(19): 10873–10879.

Gotte, B., L. Liu and G. M. McInerney (2018). “The Enigmatic Alphavirus Non-Structural Protein 3 (nsP3) Revealing Its Secrets at Last.” Viruses 10(3).

Hasanifard, L., R. Sheervalilou, M. Majidinia and B. Yousefi (2019). “New insights into the roles and regulation of SphK2 as a therapeutic target in cancer chemoresistance.” J Cell Physiol 234(6): 8162–8181.

Igarashi, N., T. Okada, S. Hayashi, T. Fujita, S. Jahangeer and S. Nakamura (2003). “Sphingosine kinase 2 is a nuclear protein and inhibits DNA synthesis.” J Biol Chem 278(47): 46832–46839.

Kaariainen, L. and T. Ahola (2002). “Functions of alphavirus nonstructural proteins in RNA replication.” Prog Nucleic Acid Res Mol Biol 71: 187–222.

Kennedy, A. C., J. Fleming and L. Solomon (1980). “Chikungunya viral arthropathy: a clinical description.” J Rheumatol 7(2): 231–236.

Kihara, A., Y. Anada and Y. Igarashi (2006). “Mouse sphingosine kinase isoforms SPHK1a and SPHK1b differ in enzymatic traits including stability, localization, modification, and oligomerization.” J Biol Chem 281(7): 4532–4539.

Kim, D. Y., J. M. Reynaud, A. Rasalouskaya, I. Akhrymuk, J. A. Mobley, I. Frolov and E. I. Frolova (2016). “New World and Old World Alphaviruses Have Evolved to Exploit Different Components of Stress Granules, FXR and G3BP Proteins, for Assembly of Viral Replication Complexes.” PLoS Pathog 12(8): e1005810.

Lark, T., F. Keck and A. Narayanan (2017). “Interactions of Alphavirus nsP3 Protein with Host Proteins.” Front Microbiol 8: 2652.

LaStarza, M. W., J. A. Lemm and C. M. Rice (1994). “Genetic analysis of the nsP3 region of Sindbis virus: evidence for roles in minus-strand and subgenomic RNA synthesis.” J Virol 68(9): 5781–5791.

Malet, H., B. Coutard, S. Jamal, H. Dutartre, N. Papageorgiou, M. Neuvonen, T. Ahola, N. Forrester, E. A. Gould, D. Lafitte, F. Ferron, J. Lescar, A. E. Gorbalenya, X. de Lamballerie and B. Canard (2009). “The crystal structures of Chikungunya and Venezuelan equine encephalitis virus nsP3 macro domains define a conserved adenosine binding pocket.” J Virol 83(13): 6534–6545.

McPherson, R. L., R. Abraham, E. Sreekumar, S. E. Ong, S. J. Cheng, V. K. Baxter, H. A. Kistemaker, D. V. Filippov, D. E. Griffin and A. K. Leung (2017). “ADP-ribosylhydrolase activity of Chikungunya virus macrodomain is critical for virus replication and virulence.” Proc Natl Acad Sci U S A 114(7): 1666–1671.

Meshram, C. D., P. Agback, N. Shiliaev, N. Urakova, J. A. Mobley, T. Agback, E. I. Frolova and I. Frolov (2018). “Multiple Host Factors Interact with the Hypervariable Domain of Chikungunya Virus nsP3 and Determine Viral Replication in Cell-Specific Mode.” J Virol 92(16).

Morchang, A., R. C. H. Lee, P. T. Yenchitsomanus, G. P. Sreekanth, S. Noisakran, J. J. H. Chu and T. Limjindaporn (2017). “RNAi screen reveals a role of SPHK2 in dengue virus-mediated apoptosis in hepatic cell lines.” PLoS One 12(11): e0188121.

Mutso, M., A. M. Morro, C. Smedberg, S. Kasvandik, M. Aquilimeba, M. Teppor, L. Tarve, A. Lulla, V. Lulla, S. Saul, B. Thaa, G. M. McInerney, A. Merits and M. Varjak (2018). “Mutation of CD2AP and SH3KBP1 Binding Motif in Alphavirus nsP3 Hypervariable Domain Results in Attenuated Virus.” Viruses 10(5).

Neuvonen, M. and T. Ahola (2009). “Differential activities of cellular and viral macro domain proteins in binding of ADP-ribose metabolites.” J Mol Biol 385(1): 212–225.

Neuvonen, M., A. Kazlauskas, M. Martikainen, A. Hinkkanen, T. Ahola and K. Saksela (2011). “SH3 domain-mediated recruitment of host cell amphiphysins by alphavirus nsP3 promotes viral RNA replication.” PLoS Pathog 7(11): e1002383.

Pialoux, G., B. A. Gauzere, S. Jaureguiberry and M. Strobel (2007). “Chikungunya, an epidemic arbovirosis.” Lancet Infect Dis 7(5): 319–327.

Pohjala, L., A. Utt, M. Varjak, A. Lulla, A. Merits, T. Ahola and P. Tammela (2011). “Inhibitors of alphavirus entry and replication identified with a stable Chikungunya replicon cell line and virus-based assays.” PLoS One 6(12): e28923.

Reid, S. P., S. R. Tritsch, K. Kota, C. Y. Chiang, L. Dong, T. Kenny, E. E. Brueggemann, M. D. Ward, L. H. Cazares and S. Bavari (2015). “Sphingosine kinase 2 is a chikungunya virus host factor co-localized with the viral replication complex.” Emerg Microbes Infect 4(10): e61.

Schilte, C., F. Staikowsky, T. Couderc, Y. Madec, F. Carpentier, S. Kassab, M. L. Albert, M. Lecuit and A. Michault (2013). “Chikungunya virus-associated long-term arthralgia: a 36-month prospective longitudinal study.” PLoS Negl Trop Dis 7(3): e2137.

Shida, D., K. Takabe, D. Kapitonov, S. Milstien and S. Spiegel (2008). “Targeting SphK1 as a new strategy against cancer.” Curr Drug Targets 9(8): 662–673.

Shin, G., S. A. Yost, M. T. Miller, E. J. Elrod, A. Grakoui and J. Marcotrigiano (2012). “Structural and functional insights into alphavirus polyprotein processing and pathogenesis.” Proc Natl Acad Sci U S A 109(41): 16534–16539.

Simon, F., E. Javelle, M. Oliver, I. Leparc-Goffart and C. Marimoutou (2011). “Chikungunya virus infection.” Curr Infect Dis Rep 13(3): 218–228.

Yamane, D., D. R. McGivern, E. Wauthier, M. Yi, V. J. Madden, C. Welsch, I. Antes, Y. Wen, P. E. Chugh, C. E. McGee, D. G. Widman, I. Misumi, S. Bandyopadhyay, S. Kim, T. Shimakami, T. Oikawa, J. K. Whitmire, M. T. Heise, D. P. Dittmer, C. C. Kao, S. M. Pitson, A. H. Merrill, Jr., L. M. Reid and S. M. Lemon (2014). “Regulation of the hepatitis C virus RNA replicase by endogenous lipid peroxidation.” Nat Med 20(8): 927–935.

